# From a different angle: genetic diversity underlies differentiation of waterlogging-induced epinasty in tomato

**DOI:** 10.1101/2022.12.02.518852

**Authors:** B. Geldhof, J. Pattyn, B. Van de Poel

## Abstract

In tomato, downward leaf bending is a morphological adaptation towards waterlogging, which has been shown to induce a range of metabolic and hormonal changes. This kind of functional trait is often the result of a complex interplay of regulatory processes starting at the gene level, gated through a plethora of signaling cascades and modulated by environmental cues. Through phenotypical screening of a population of 54 tomato accessions in a Genome Wide Association Study (GWAS), we have identified target genes potentially involved in plant growth and survival during waterlogging and subsequent recovery. Changes in both plant growth rate and epinastic descriptors revealed several associations to genes possibly supporting metabolic activity in low oxygen conditions in the root zone. In addition to this general reprogramming, some of the targets were specifically associated to leaf angle dynamics, indicating these genes might play a role in the induction, maintenance or recovery of differential petiole elongation in tomato during waterlogging.

## Introduction

Leaf angle is an important agricultural trait in both monocot and dicot crops that ensures optimal light interception and resource allocation for plant growth and development. Especially for cereal crops, growing in high densities, this feature has gained a lot of attention as a possible target for yield optimization (Mantilla-Perez et al., 2020; Mantilla-Perez & Salas Fernandez, 2017). Leaf angle is, however, a dynamic trait, and plants can adjust their leaf posture to enhance survival in suboptimal conditions. Arabidopsis, for example, wields the hyponastic response as a strategy to overcome low-oxygen stress (Hebelstrup et al., 2012; Rauf et al., 2013), compete with neighbors by means of the shade-avoidance response (Pantazopoulou et al., 2017) and improve heat dissipation under high temperatures (Park et al., 2019). This upwards bending is the outcome of an array of hormonal cues that control leaf angle in space and time. Several key regulators, transducing hormonal signals through genetic programming, have been described, revealing a common framework for posture control during abiotic stress in Arabidopsis.

Many of the known regulators of hyponasty are involved in hormonal signaling. During waterlogging in Arabidopsis, hyponastic growth is triggered by SPEEDY HYPONASTIC GROWTH (SHYG), which acts upstream of 1-AMINOCYCLOPROPANE-1-CARBOXYLIC ACID (ACC) OXIDASE5 (ACO5), a key enzyme in ethylene biosynthesis (Rauf et al., 2013). Ethylene induced hyponasty itself is further controlled by reorientation of cortical microtubules, leading to differential elongation in the petiole (Polko et al., 2012). This process is likely further regulated by brassinosteroid (BR) signaling (Polko et al., 2013). Concomitantly, key photomorphogenic transcription factors are activated to, independent of light signaling, induce upward leaf movement in *Rumex palustris*, indicating shade avoidance and low-oxygen responses rely on a shared arsenal of regulators (van Veen et al., 2013).

In contrast to Arabidopsis wielding hyponasty as a stress response, other species such as tomato, actively bend their leaves downwards during waterlogging, which is called epinasty. It is likely that the epinastic response is also the output of a series of hormone-regulated processes, starting with the accumulation of ACC in the hypoxic roots (Olson et al., 1995; Shiu et al., 1998). ACC is then transported to the shoot (English et al., 1995; Jackson & Campbell, 1976) and converted into ethylene (English et al., 1995), which converges on AUXIN/INDOLE-3-ACETIC ACID 3 (SlIAA3), an integrator of ethylene and auxin responses that is partially controlling leaf epinasty (Chaabouni et al., 2009a). As a result, differential elongation is likely, but not exclusively, gated by auxin redistribution (Lee et al., 2008). *SlIAA3* expression is also modulated by BR, as impaired functioning of DWARF (DWF), a BR biosynthesis gene, represses both ethylene and BR production, strongly induces *SlIAA3* expression (in the stem) and evokes a hyponastic phenotype (Li et al., 2016).

In contrast to the genetic networks resulting in asymmetric petiole growth during hyponasty in Arabidopsis, little is known about the central nodes of such a regulatory network in waterlogging-induced epinasty in tomato. In search for these regulators, we performed a multi-trait and longitudinal (time-series) Genome-Wide Association Study (GWAS) in tomato, revealing that leaf posture control is a highly dynamic trait controlled by many factors. This study shows that tomato harbors genetic diversity with respect to leaf epinasty as an adaptive strategy to survive waterlogging.

## Materials and methods

### Plant material and growth conditions

Tomato seeds (*Solanum lycopersicum*) of 54 accessions, part of a larger collection described in Tieman et al. (2017), were obtained from the Tomato Genetics Resource Center (TGRC) or were kindly provided by Prof. D. Zamir. Seeds were germinated in soil and afterwards transferred to rockwool blocks. Tomato seedlings were grown in the greenhouse until they reached approximately the eighth leaf stage, after which they were used for experiments. In the greenhouse, temperature was set at 18 °C (day and night) with a relative humidity between 65 – 70 %. Additional illumination (SON-T) was provided if solar light intensity dropped below 250 W m^-2^. The plants received fertigation through automated drip irrigation.

### Waterlogging treatment and phenotyping

At the eighth leaf stage, half of the plants of each accession was transferred to individual containers and subjected to a three-day waterlogging treatment. The container was filled with water up to 4 cm above the rockwool surface. Oxygen depletion during waterlogging was achieved by natural root respiration, leading to rapid hypoxia, generally within 3 – 8 h (Geldhof et al., 2021). Afterwards, the root system was drained and reoxygenated and plants were monitored for another three days. At the start of the treatment and during reoxygenation, plant height was measured daily. Leaf angle dynamics were monitored using real-time leaf angle sensors (Geldhof et al., 2021), attached to the petiole of the first eight leaves to take into account ontogenic differences. Four out of the 54 accessions were not shared for growth and leaf angle phenotyping (52 unique accessions per dataset).

### Genome-wide association study

Time-series continuous leaf angle data were transformed into key traits describing leaf angle dynamics (see results and Supplemental Table S1). GWA mapping was performed based on the study by Tieman et al. (2017), using the EMMAX algorithm (with missingness < 10 % and maf > 5 %) (Kang et al., 2010). Monoallelic SNP files were filtered, only retaining the accessions used in this study, and converted into biallelic and binary format using PLINK. The significant P-value threshold was determined using the Genetic type 1 Error Calculator (GEC). The power of the analysis was verified using randomized phenotypes and potential environmental interactions were tested with the Genome-wide Efficient Mixed Model Association (GEMMA) algorithm (Zhou & Stephens, 2014). Design parameters for the model (genotype – phenotype combinations) were bash scripted in R (R Core Team, 2019) and the analysis was run for each of the above phenotypical variables and for the leaf angle itself at 10 minute intervals.

Plant growth during waterlogging and recovery were defined as the average growth over the 3-day waterlogging period or as the slope of a linear model during recovery respectively. The ontogenic effect was incorporated in the GWAS through quadratic models with leaf age as independent and angle descriptors as dependent variables. Leaf age-specific time-series were represented using Functional Principal Component Analysis (FPCA) on downsampled series, angular descriptors and their Best Linear Unbiased Estimators (BLUE). For each of these variables, average values were determined per treatment and their differences were fed to the EMMAX algorithm.

### RNA extraction and RT-qPCR

Tomato petioles were sampled and adaxial and abaxial sections were snap frozen and ground in liquid nitrogen. RNA was extracted using the GeneJET Plant RNA Purification Mini Kit protocol (Thermo Scientific). DNA was removed using the RapidOut DNA Removal Kit (Thermo Scientific). Total RNA yield was assessed on the Nanodrop (Nanodrop Technology) and RNA quality was verified through gel electrophoresis. cDNA was synthesized using the iScript cDNA Synthesis Kit protocol (BIO-RAD). RT-qPCR was performed with a StepOnePlus (Applied Biosystems) for 40 cycles (SsoAdvanced Universal SYBR Green Supermix; Bio-Rad). Relative quantification and PCR efficiency was calculated using a standard dilution series for each primer-pair. Four reference genes (EF1a, PP2AC, RPL2 and TIP41) were selected based on earlier findings (Van de Poel et al., 2012) for normalization. Primers are listed in Supplemental Table S2.

### Data analysis

All analyses and visualizations were performed in R. Outliers of phenotypical variables were removed prior to the GWAS. Functional Principal Component Analysis (FPCA) was carried out with the *fdapace* package (Zhou et al., 2022). Gene expression levels were compared between control and waterlogging treatment with a Wilcoxon test (α = 0.05). Data visualization was done with the *ggplot* package (Wickham & Winston, 2019).

## Results

### Growth dynamics during waterlogging reveals an accession-dependent pause-recovery strategy

Waterlogging results in suboptimal conditions eventually leading to reduced growth. We quantified this growth effect for 52 tomato accessions after 3 days of waterlogging (Figure 1A & B) and during the subsequent 3-day recovery phase (Figure 1C & Supplemental Figure S1) by measuring changes in plant height. While most of these accessions had a reduced growth rate, some of them retained a more steady growth during waterlogging (e.g. accession 53). Other accessions showed a growth pause during waterlogging, followed by enhanced growth during recovery (e.g. accessions 10, 13, 38, below the identity line in Figure 1C) or seemed unaffected in both conditions (e.g. accessions 21, 43, 47). The extent of this growth reduction was in part determined by the initial plant size, indicating that plant development guides waterlogging responses of tomato (significant effect; Supplemental Figure S2). Overall, the outgrowth of new leaves was delayed for all accessions, but with differences between accessions (Figure 1D). The effect of waterlogging and reoxygenation is clearly visible in the growth dynamics of certain accessions (exemplified in Figure 1E for the accessions below the identity line in Figure 1C).

**Figure 1:**
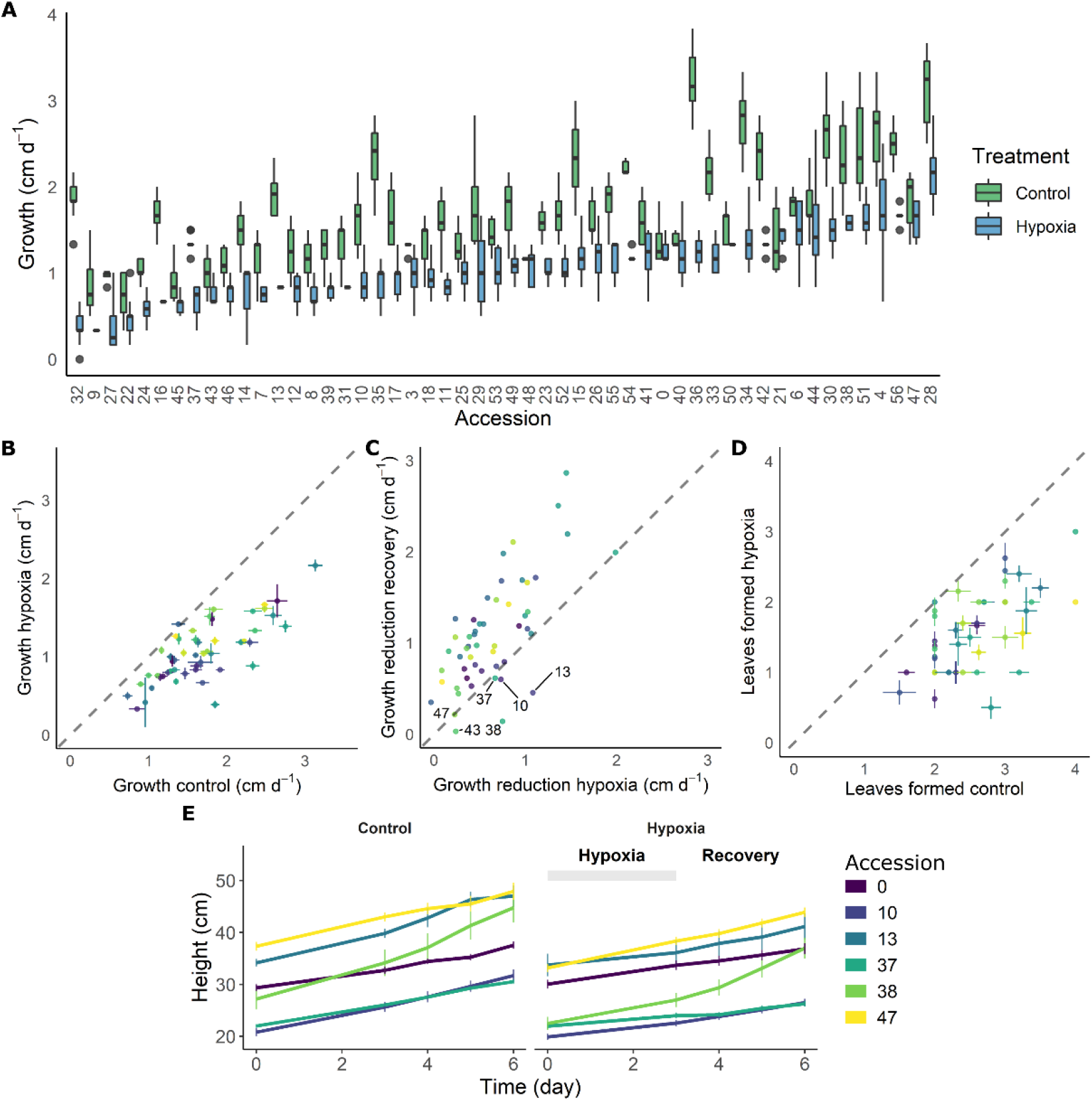
Effect of 3 days of waterlogging and recovery on growth of 52 different tomato accessions. (A) Natural variation in growth rate under control conditions and during waterlogging (n = 52). (B – C) Shift in average growth rate of the accessions during (B) 3 days of waterlogging and (C) subsequent 3 days of recovery. (D) Effect of waterlogging and recovery on leaf emergence for the different accessions. (E) Growth curves of 6 accessions during waterlogging and recovery as compared to control conditions. Vertical and horizontal bars in (A, C, E) are 95 % confidence interval.

### Growth during waterlogging and recovery is genetically regulated

The genetic variability in growth reduction and reinitiation indicates that different accessions wield different strategies to overcome waterlogging. We next investigated if variability of this resilience or recovery potential is associated with single nucleotide polymorphisms (SNPs). Therefore, both differential growth over the entire waterlogging (*G*_*C*_ *– G*_*H*_; C = control, H = hypoxia) and subsequent recovery (*G*_*C*_ *– G*_*R*_; C = control, R = recovery) treatment were used as input phenotypes for a GWAS analysis. As the 52 accessions used in this study originated from a larger population of 402 accessions (described in Tieman et al. (2017)), we decided to incorporate the overarching population genetic variability in the analysis. To this extent, PCA scores were derived based on the SNP data for the entire population and the PCs of the 52 accessions were retained as covariates in the analysis. This methodology allowed us to make full use of the underlying population structure (Figure 2A), largely covered by the 52 accessions, except for a subgroup of divergent heirloom varieties.

**Figure 2:**
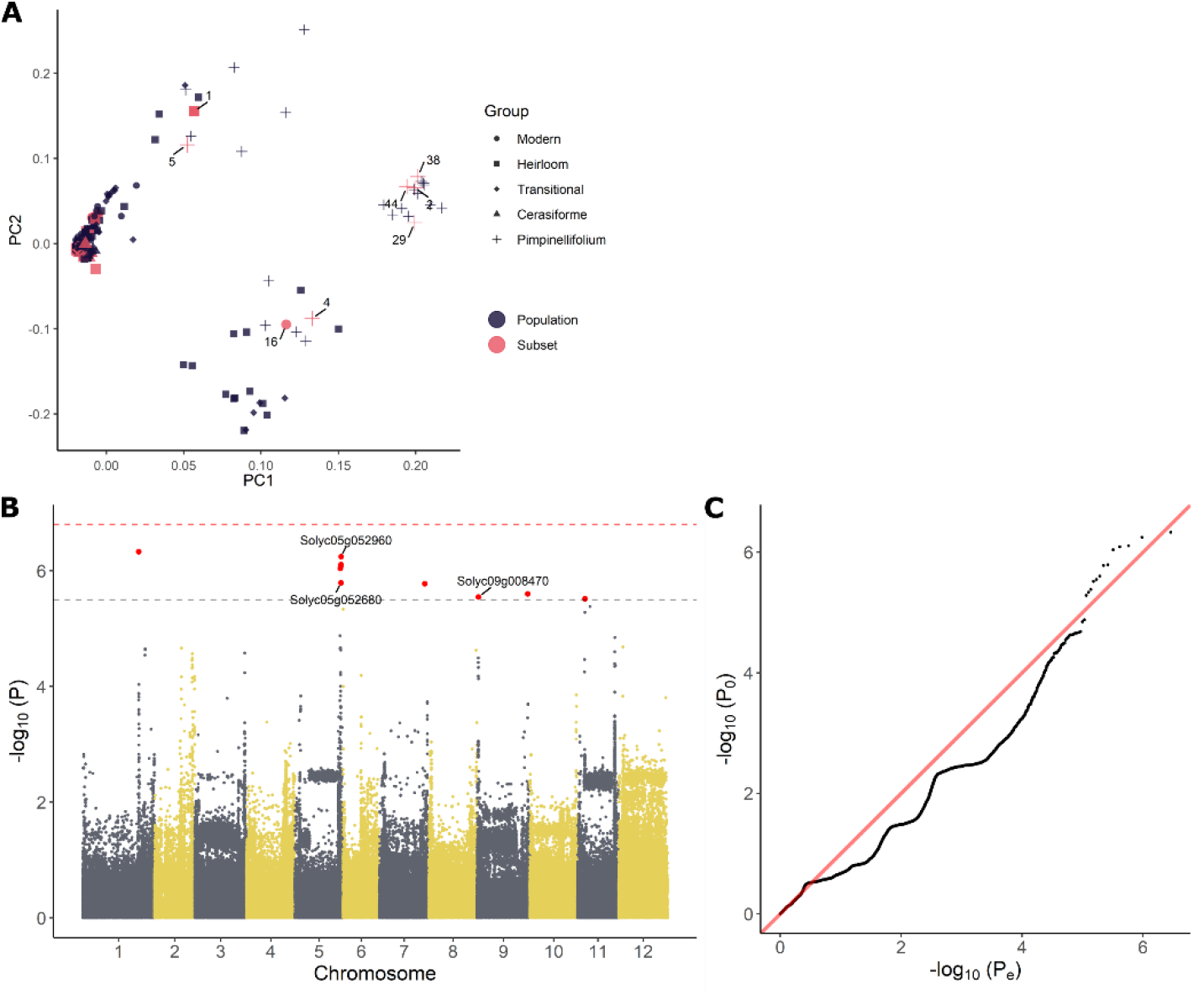
GWAS analysis on growth responses during waterlogging of tomato. (A) Population structure of the 54 tomato accessions subset used in this study (red) and those of the full population (402 accessions) described in Tieman et al. (2017) (black). (B) Manhattan and (B) QQplot of the GWAS with differential growth during waterlogging as phenotype. Dashed gray and red lines in (C) indicate suggestive and significant -log_10_(P) values (-log_10_(P_sug_) = 5.50 and -log_10_(P_sign_) = 6.80). SNPs exceeding the suggestive threshold are colored red. Annotated genes are labeled with their corresponding Solyc ID.

The results of the GWAS were validated based on the Manhattan and QQplot plot (Figure 2B – C). Despite deflation (λ = 0.907 & λ = 0.894), the analysis picked up targets exceeding both the suggestive and significant –log_10_(P) values (P_sug_ = 5.50 and P_sign_ = 6.80), indicating these SNPs could be associated with growth regulation during waterlogging. The SNPs that could be annotated were retained and are listed in (Supplemental Table S3). Two of these SNPs reoccurred during both waterlogging and recovery. The first SNP was located in the 3’ UTR region of the gene *BTB/POZ domain containing protein expressed* (Solyc05g052960), a homolog of the *BTB/POZ and MATH domain-containing protein 3* (*BPM3*) gene in Arabidopsis (AT2G39760), known to target HOMEOBOX PROTEIN 6 (AtHB6) to regulate growth and ABA signaling (Lechner et al., 2011). The second SNP was located in an exon of a BAHD acyltransferase DCR/HXXXD-type acyl-transferase/hydroxycinnamoyl transferase family protein gene (Solyc05g052680; ortholog of AT2G39980), possibly involved in specialized metabolism.

### Waterlogging-induced epinasty as a conserved trait in tomato

The above analysis revealed that growth during waterlogging is genetically determined, leading to different survival strategies. One strategy is to maintain growth during waterlogging and recovery, while another is the induction of a pause phase, probably encompassing morphological adaptations such as the formation of aerenchyma and/or adventitious roots. Another morphological adaptation towards root hypoxia is leaf epinasty. To explore the genotype-specific variability in angular dynamics during waterlogging, we monitored leaf movements with digital sensors (Geldhof et al., 2021), taking into account leaf age. High resolution time-series data were compressed in 14 different traits and their derivative variables (e.g. best linear unbiased estimators (BLUEs)) describing the epinastic response (Figure 3A – B), including total angular change during hypoxia and the maximal rate of change during the entire sequence (hypoxia and recovery). We first verified the usability of these descriptors by PCA, demonstrating that they could distinguish phenotypic variation during the treatment (Figure 3C – D). On the other hand, many of the traits were grouped, indicating they were inherently correlated. For example, the total angular change (θ_end_ – θ_0_) and rate of change during hypoxia (Δ(θ)_max_) were associated, meaning that fast epinastic responses and large epinastic curvatures go hand in hand. Interestingly, this does not necessarily imply that these plants had a strong recovery potential after the treatment.

**Figure 3:**
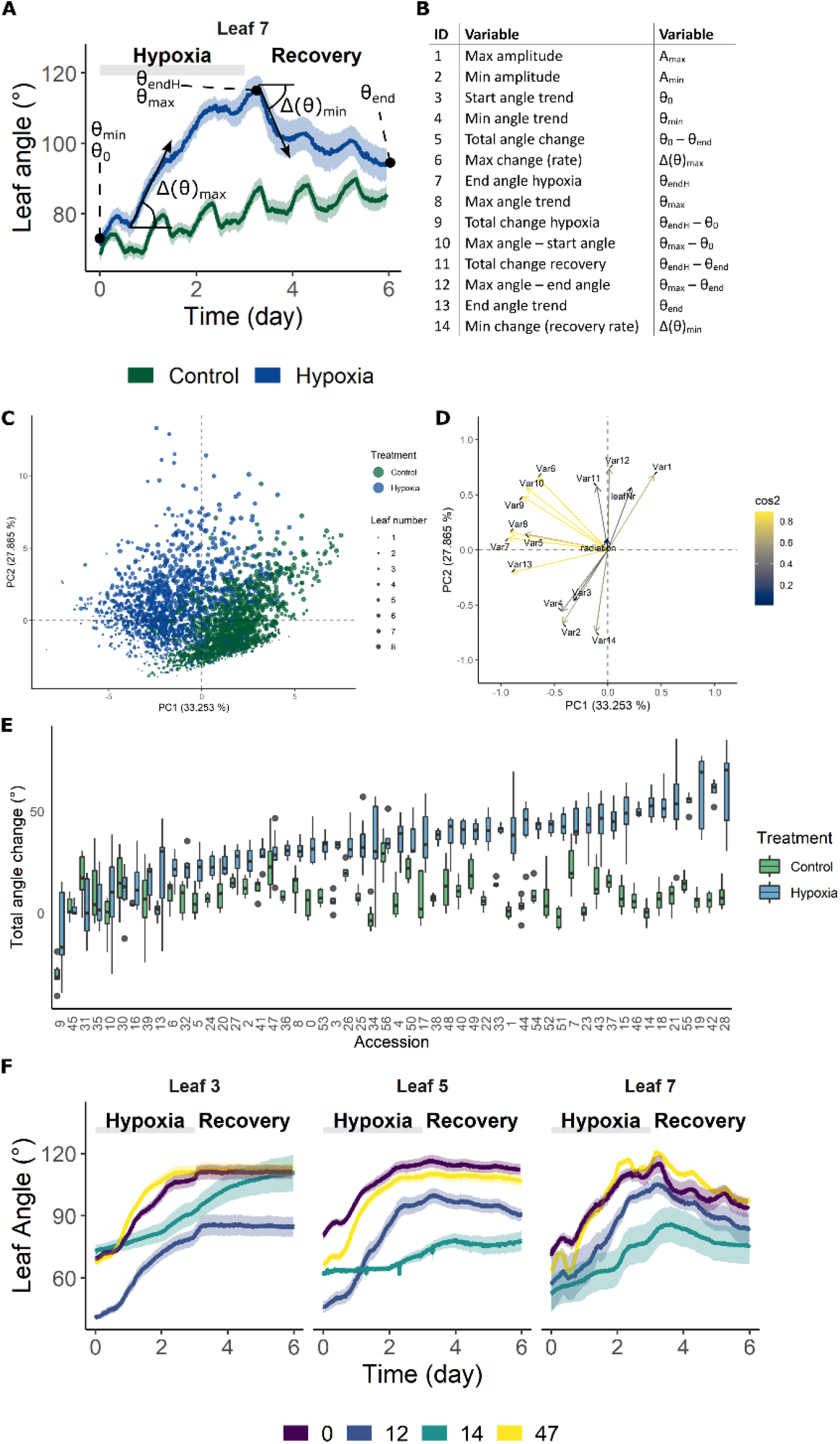
Phenotypic variation of leaf angle traits during waterlogging and recovery of tomato. (A – B) Visualization of the 14 angle descriptors used in this study. (C – D) PCA on the angle descriptors showing the effect of waterlogging (treatment) and leaf development (leaf number) on leaf posture. (C) Observation plot of individual leaves and (D) variable plot of the descriptors defined in (A – B) with their quality of representation. (E) Natural variation in total leaf angle change (θ_end_ – θ_0_) in control conditions and after 3 days of waterlogging and subsequent recovery of leaf 5. (F) Leaf angle dynamics of leaves of different ages (leaf 3, 5 and 7) of 4 different accessions. Colored bands represent the 95 % confidence interval.

The above descriptors are not only capable of describing waterlogging-induced changes in leaf posture, but they also vary between different accessions (Figure 3E – F), providing a phenotypical fingerprint for the GWAS. An example of the effect of waterlogging on total leaf angle change and related leaf posture traits for leaf 5 of 52 accessions is given in Figure 3E and Supplemental Figure S3 respectively. While some of the accessions show major leaf repositioning throughout waterlogging and recovery (e.g. 14, 18, 42), others seem to recover (e.g. 10, 30) (Figure 3E). By combining all of these angular variables (Figure 3B), an epinastic heatmap of the waterlogging effect could be devised for each of the accessions (Supplemental Figure S3). This genetic variability, leading to distinctive dynamic leaf movements, is exemplified for different accessions (Figure 3F) and developmental stages (leaf 3, 5 and 7), indicating that both genotype and ontogeny guide leaf movement during waterlogging and recovery.

### Angular dynamics during waterlogging is associated with distinct SNP signatures

Similarities in leaf posture and movements between different tomato accessions indicate that epinasty is a conserved response towards waterlogging stress in this species. To investigate the genetic basis of this mechanism, each of the angular traits was used separately as a phenotype for GWAS analysis, taking into account ontogeny. Besides these angular descriptors, we also fed the model with time-series angle data, revealing a number of critical points in the regulation of the epinastic response. At certain time points, both the count of associated SNPs and the rate of angular change induced by waterlogging were strongly elevated (Figure 4A – B). Both profiles also indicate that this regulation might be gated by the circadian clock, with major shifts caused by waterlogging starting mostly in the morning, following the oscillatory leaf movement. By analyzing the timing of divergence between waterlogging-treated plants and control plants, we quantified the onset of the epinastic response in Ailsa Craig, as a reference threshold for early responses. After 400 – 500 minutes (6 – 8 h), the rate of angular change increased significantly in leaves of waterlogged Ailsa Craig plants, providing a good reference time point for early epinastic responses in the GWAS.

**Figure 4:**
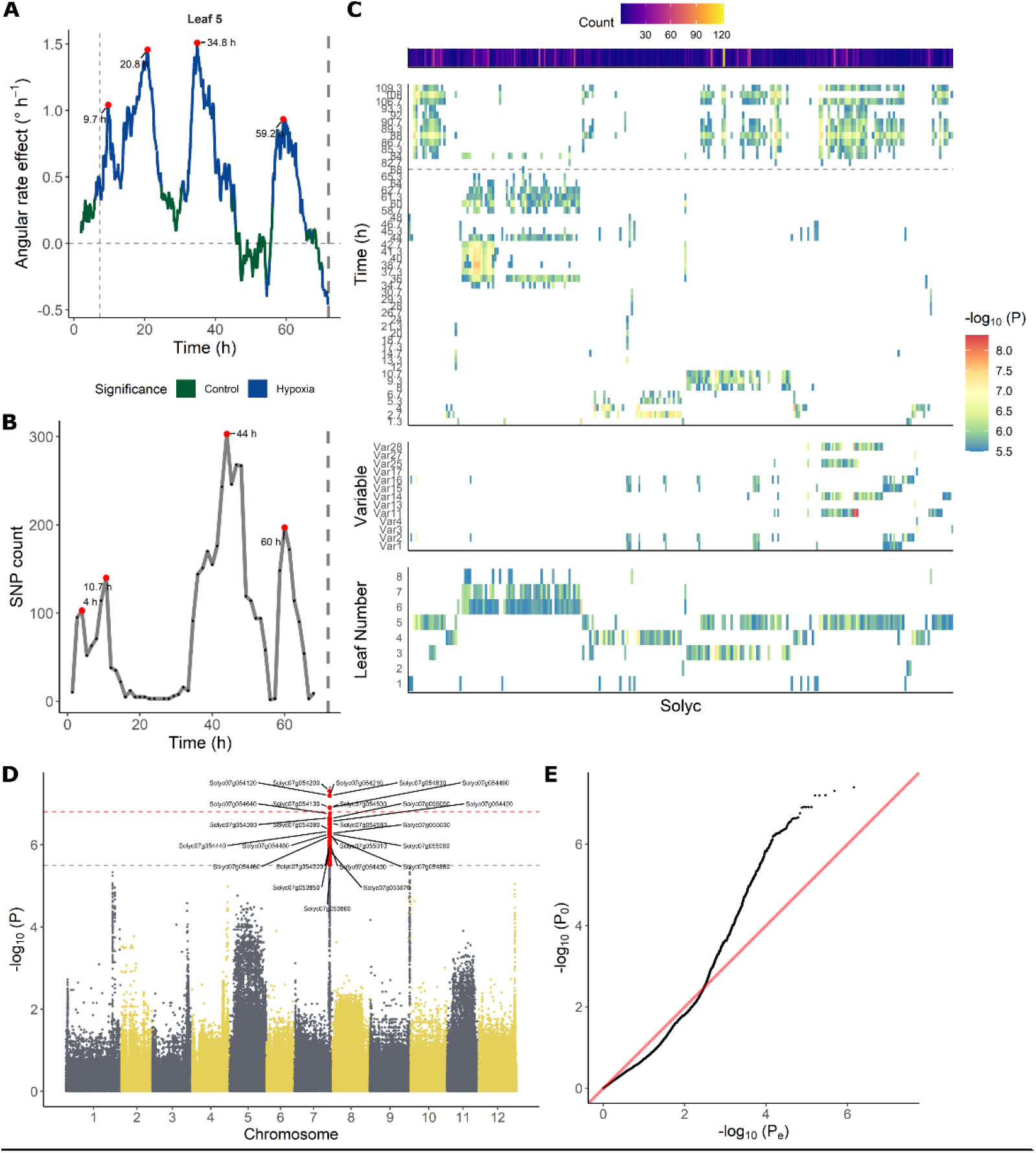
Association between leaf angle dynamics and SNP data during waterlogging and recovery in tomato. (A – B) Representation of (A) the effect of waterlogging on rate of angular change in leaf 5 and (B) SNP count, highlighting (red dots) certain periods of higher angular activity and higher number of associated SNPs. (C) Ontogenic (leaf 1 – 8) and time-dependent profile of annotated genes (Solyc number) associated with leaf angle changes during waterlogging. The heat maps depict the occurrence of significant associations of a certain gene in time (upper), with an angle variable (middle) and with leaf age (lower). (D – E) Manhattan and (D) QQplot of the GWAS analysis with leaf angle around a pivotal point (10.7 h) as phenotype (leaf 3). Dashed gray and red lines in (D) indicate suggestive and significant -log_10_(P) values (-log_10_(P_sug_) = 5.50 and -log_10_(P_sign_) = 6.80). SNPs exceeding the suggestive threshold are colored red. Annotated genes are labeled with their corresponding Solyc ID.

Next, we filtered the SNP targets, retaining only SNPs with a –log_10_(P) value above the suggestive P-value and located in non-inflated chromosomic regions. These regions were determined by PCA on genome-wide P-test statistics, only retaining chromosomic regions with a limited total count of SNPs and –log_10_(P) (non-outliers in 5 subsequent PCAs). Only SNPs that could be annotated were further analyzed (Supplemental Table S4), revealing a grouping structure based on leaf age (lower panel Figure 4C) and angle dynamics (upper panel Figure 4C). SNPs associated with fast responses (< 10 h) were more enriched in mature leaves (leaf 3 – 5), and some of these SNPs were also detected during recovery (> 72 h). SNPs associated with angular responses in young leaves were mostly detected during the later stages of the treatment (> 34 h), corresponding to the second period of maximal divergence of angular rate between waterlogging and control (Figure 4A).

### Specific gene groups are associated with early and late angular responses

In general, several SNPs and genes reoccurred throughout GWAS analyses for different phenotypes, and were associated with similar variables or subsequent time points in the time-series analysis. Given this clear time-dependent SNP profile (Figure 4B – C), we decided to tie SNP occurrence to distinct phases in the epinastic response (early: < 12 h; late: > 34 and < 72 h; recovery phase). While no genes harboring significant SNPs (higher than significant -log_10_(P)) were shared between these three phases, this was the case when including suggestive SNPs, indicating certain processes might still be shared (Supplemental Figure S4; Supplemental Table S5 – S9). To gain more insight in the early angle response, we focused on SNPs that were only detected at the onset of the waterlogging treatment (< 12 h), possibly including key activators of epinasty. Most genes harboring one or more of these SNPs also reoccurred at later time-points and were mainly associated with responses for leaf 3 and 4. A small-scale GO analysis showed that genes involved in plasma membrane processes (9), defense responses (4), signal transduction (4) and two vacuolar (2 + 3) and RNA processing (3 + 2) categories were relatively abundant. Furthermore, a group of protein serine/threonine kinases and receptor like kinases was well-represented in this set (5).

We further investigated genes with SNPs having a –log_10_(P-value) above the significance threshold and associated with early angle changes (< 12 h) (24 genes; Supplemental Table S5). The most reoccurring genes harboring SNPs were Cc-nbs-lrr, resistance protein (Solyc04g007050), Genomic DNA chromosome 3 P1 clone MYA6 (Solyc03g120490), S-receptor kinase-like protein 1 (Solyc07g005110), Vacuolar ATPase subunit H protein (Solyc07g005940), Ribonuclease 3-like protein 3 (Solyc07g005030) and Response regulator 23 (RR23) (Solyc07g005140). Other significant targets were involved in RNA processing, transport (e.g. ABC transporter C family member 2 (ABCC2)) and energy metabolism (e.g. Aldehyde dehydrogenase (ALDH)).

Besides these genes with SNPs specifically associated with early angular responses, other early-response genes reoccurred during the reoxygenation phase (when using suggestive SNPs), indicating they might also be involved in leaf repositioning. Genes discovered within the early waterlogging and recovery phase included a LRR receptor-like serine/threonine-protein kinase (EMS1-like, Solyc07g054120, 76 SNPs), Receptor-like kinase (SRF4, Solyc07g054500, 11 SNPs), a gene involved in energy metabolism (ATP synthase I-like, Solyc07g055050, 17 SNPs) and an unknown protein (Solyc07g054130, 22 SNPs). Several of these SNPs were associated with both early responses in older leaves (leaf 3) and late responses in younger leaves (leaf 5), indicating the corresponding genes are ontogenically regulated in time. Genes present in both of these subsets included Transmembrane protein 34 (Solyc07g054320, 142 SNPs), B3 domain-containing protein (homolog of RELATED TO VERNALIZATION1 (AtRTV1), Solyc07g054630, 48 SNPs) and a NADH dehydrogenase (Solyc01g088350, 40 SNPs).

Leaf movement in young leaves is not limited to repositioning during reoxygenation. A set of 3 genes was specifically and significantly associated with movement in young leaves, mostly around 36 h after the start of the waterlogging treatment. This group consisted of a Polyamine oxidase (PO4, Solyc03g031880, 45 SNPs), a Coatomer protein epsilon subunit family protein (Solyc02g069590, 22 SNPs), and a LOB domain protein 4 (LBD4, Solyc02g069440, 10 SNPs).

Another 37 genes were present both during the late response (> 34 h) and during subsequent recovery, indicating they might be involved in late epinastic bending or stress recovery. There was no clear grouping of these individual genes, suggesting they might regulate distinct processes. However, multiple genes were involved in nuclear processing (6), including light-dependent short hypocotyls 1 (Solyc02g069510, 13 SNPs), a histone deacetylase (Solyc09g091440 22 SNPs) and a MYB85 transcription factor (Solyc07g054980, 10 SNPs). Also, a large number of SNPs exceeded the suggestive P-value, almost reaching the significant threshold, including Homeobox-leucine zipper protein AtHB14 (Solyc03g031760, 9 SNPs) and Auxin response factor 1-2 (ARF8B, Solyc03g031970, 7 SNPs).

A group of SNPs specifically associated with the recovery phase located to genes mainly enriched in membrane-bound receptor kinases. Besides these signaling components, the analysis also picked up Glucose transporter 8 (Solyc12g089180) during the reoxygenation phase. Other targets were mainly involved in ATP (8) and protein binding (6) and in nuclear processes (7), including histone modification (Solyc02g079300, Solyc09g091440 and Solyc01g008120) and transcription (e.g. Histidine kinase/DNA gyrase B (Solyc09g092100), MYB85 (Solyc07g054980)). This set also contained another aldehyde oxidase gene (AO4 pseudogene, Solyc01g088170, 66 SNPs) and two genes involved in specialized metabolism (Cinnamoyl CoA reductase-like protein (PAR1, Solyc01g008530) and Hydroxycinnamoyl-CoA shikimate/quinate hydroxycinnamoyl transferase (Solyc02g079490)).

### Identification of potential regulatory targets

Next, we selected a few candidate target genes based on their timing and possible developmental association, to further study their involvement in leaf angle regulation (Table 1). Therefore, we verified if these genes were asymmetrically expressed within the petiole during waterlogging. Their gene expression level was quantified in both the abaxial and adaxial side of the petiole of leaf 5 after 48 h of waterlogging (Figure 5). Most of the selected targets showed only mild but non-significant changes. The lack of differences between control and epinastic petioles, is possibly due to sampling constraints and sample characteristics (e.g. containing a relative large proportion of vascular tissue). On the other hand, one of the several receptor-like kinases associated with the epinastic response at different time points (EMS-like1) did show a major shift during waterlogging. This LRR receptor-like serine/threonine-protein kinase, RLP (Solyc07g054120), harbored a large number of significant SNPs, especially at 560 minutes (9.33 h) after the start of the treatment in leaf 3 and also reoccurred during recovery in leaf 5. This gene was strongly downregulated in both petiole sections after 48 h of waterlogging (Figure 5).

**Table 1:** Target gene selection for RT-qPCR

| Solyc ID | Gene ID | Alternative annotation | Timing | Leaf |
| --- | --- | --- | --- | --- |
| Solyc07g005140 | RR23 |  | Early | Mature (4) |
| Solyc07g054120 | LRR receptor-like serine/threonine-protein kinase, RLP | EMS-like 1 | Early<br>Recovery | Old (3)<br>Mature (5) |
| Solyc02g069440 | LBD4 |  | Late | Young (6 – 7) |
| Solyc02g079400 | Nitric oxide synthase interacting protein (NOSIP) | CSU1 | Recovery | Old – Young |
| Solyc07g054500 | Receptor like kinase, RLK | SRF3/4 | Early<br>Recovery | Old (3)<br>Mature (5) |

**Figure 5:**
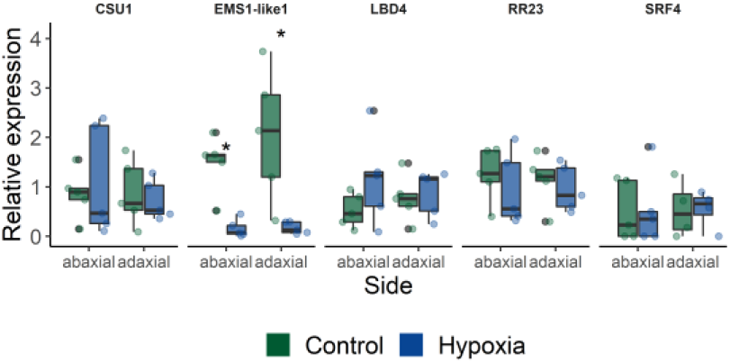
Expression of a subset of target genes identified in the GWAS (CSU1 = COP1 SUPPRESSOR 1; EMS1-like 1 = EXCESS MICROSPOROCYTES1-LIKE 1; LBD4 = LOB DOMAIN PROTEIN 4; RR23 = RESPONSE REGULATOR 23; SRF4 = STRUBBELIG-receptor family 4). Gene expression was quantified in abaxial and adaxial petiole sections (leaf 5) of plants subjected to waterlogging for 48 h (n = 5). Statistical significant differences (P < 0.05) are indicated with an asterisk.

## Discussion

### Waterlogging tolerance is genetically encoded in tomato

Genetic diversity has often been exploited as a natural resource for beneficial traits in breeding programs, including enhanced yield and stress resilience. For example, natural populations of Arabidopsis accessions have been screened for survival during drought (Kalladan et al., 2017) and flooding (Meng et al., 2022; Vashisht et al., 2011). However, insights in natural differentiation of waterlogging responses in crops such as tomato are mostly lacking. We have now shown that different tomato accessions display distinct growth responses when opposed to waterlogging and identified accessions with an enhanced recovery potential (Figure 1). We hypothesize that there are several survival strategies based on waterlogging resilience and tolerance. Some accessions wield a pause-strategy and display stagnant growth during waterlogging, followed by enhanced growth recovery during reoxygenation. Other accessions continue to grow during waterlogging and maintain a similar growth rate afterwards, while other accessions undergo a strong growth reduction during both phases.

Our GWAS analysis revealed that a reduced growth rate during waterlogging and recovery were associated with a limited number of genes (Figure 2), including a MATH-BTB/POZ domain containing protein (SlBTB12, Solyc05g052960). It has been shown that AtBPM3, the closest homolog of SlBTB12 in Arabidopsis, can interact with the ethylene response factor APETALA2/ERF (AP2/ERF) transcription factor family members RAP2.4b and d (Rudnik et al., 2017; Weber & Hellmann, 2009), involved in light and ethylene signaling and overall plant development (Lin et al., 2008). Through its interaction with RAP2.4, BPM3 modulates growth and drought responses, independently of ABA (Lin et al., 2008). In contrast, through degradation of AtHB6, BPM3 can also regulate plant growth and interfere with ABA signaling (Lechner et al., 2011). In tomato, expression of SlBTB12 is controlled by various hormones and environmental conditions (Li et al., 2018), indicating this transcription factor might integrate signals for growth and abiotic stress such as waterlogging.

The second gene identified in this analysis, a hydroxycinnamoyl transferase (HCT) (Solyc05g052680), is part of the phenylpropanoid pathway. The HCT clade belongs to a larger family of BAHD acyltransferases, involved in the specialized metabolism, including the production of phenolic compounds such as anthocyanin and suberin in roots (Bontpart et al., 2015; Gou et al., 2009; Molina & Kosma, 2015). In tea (*Camellia sinensis*), overexpression of one of these BAHD genes affects not only the specialized metabolism, but also plant growth (Aktar et al., 2022). How and if this *HCT* gene aids tomato plants in waterlogging resilience remains to be investigated.

### Leaf epinasty and metabolic changes are coregulated during waterlogging in tomato

The downwards bending of leaves (epinasty) reduces canopy cover, thereby limiting transpirational water losses (Else et al., 1995) and possible deleterious effects of photo-inhibition when carboxylation has ceased (Pastenes et al., 2005; Van Geest et al., 2012) during waterlogging. As such, epinasty and its recovery are part of the morphological adaptation to increase survival under low-oxygen stress conditions. We found a large genetic diversity in this waterlogging-induced epinastic bending, with some accessions showing a strong response while others displayed a mild or no response (Figure 3). The analysis also showed that shifts in angular change coincided with SNP frequency, defining key phases during waterlogging-induced epinastic bending and subsequent recovery.

Our GWAS analysis revealed a number of candidate genes that might be associated to this epinastic response, either by directly affecting leaf posture or by regulating other low-oxygen activated pathways (Figure 4). Several of these genes (e.g. Aldehyde dehydrogenase (Solyc07g005390); Amine oxidase family protein (Solyc03g031880)) were also retrieved in a transcriptomic study that assessed waterlogging of tomato (De Ollas et al., 2021), indicating they might be transcriptionally regulated during stress adaptation. Waterlogging-induced changes include rewiring of the carbohydrate and energy metabolism and the energy harvesting mechanism (Supplemental Table S10). For example, we identified SNPs in genes that include an aldehyde dehydrogenase (Solyc07g005390), beta-galactosidase (Solyc09g092160), Phosphoenolpyruvate carboxykinase (Solyc12g088160) and protochlorophyllide reductase like (Solyc07g054210). Recently, it has been shown that levels of the phytotoxic chlorophyll precursor protochlorophyllide are controlled by oxygen sensing in etiolated seedlings of tomato (Abbas et al., 2022).

Not only does this change indicate reprogramming of the energy metabolism, but also the detoxification of its ROS byproducts. Our GWAS detected SNPs in a thioredoxin reductase (Solyc03g032000), involved in ROS scavenging, and an aldehyde oxidase/xanthine dehydrogenase (AO/XDH) module potentially regulating ROS production and ABA biosynthesis (Sagi et al., 1999) (Solyc01g088200, Solyc01g088210, Solyc01g088230), indicating the need for ROS detoxification during waterlogging and recovery. Whether or not this directly affects leaf bending in tomato remains to be investigated, but XDH activity has been linked with leaf curling in Arabidopsis treated with the synthetic auxin 2,4-D (Pazmiño et al., 2014).

Besides apparent changes in the energy metabolism, our analyses also revealed some ambiguous associations related to genetic regulation, including DNA and RNA processing, small RNA synthesis and epigenetic regulation (Supplemental Table S10). This is not unexpected as plants need to activate a suite of adaptation responses to ensure their survival under waterlogging conditions. Especially epigenetic changes during stress, including chromatin remodeling, are gaining attention as they can determine the plasticity to respond adequately to changes in environmental conditions. Epigenetic changes have been observed in a range of species during flooding (Reynoso et al., 2019) and drought stress (Reynoso et al., 2022).

### Is a network of kinases involved in low-oxygen stress signaling?

The variety of morphological and metabolic changes during waterlogging are the result of fine-tuned signal transduction pathways, conveying messages within and between cells and over long distances between tissues. To our surprise, we found several SNPs in target genes involved in signaling and transport. Notably, we could identify several receptor-like kinases (RLK) and serine/threonine-protein kinases (12), including multiple G-type lectin S-receptor-like kinases (5) (Teixeira et al., 2018) and members of the wall-associated kinase (WAK) family (2) (Sun et al., 2020). Some of these targets have previously been reported to act in early and systemic signaling events in roots. For example, STRUBBELIG RECEPTOR KINASE 3 (SRF3), the closest Arabidopsis homolog to RLK (Solyc07g054500), coordinates root growth through iron sensing, while SERINE/THREONINE PROTEIN KINASE 3 (PK3/WAG1, homolog to Solyc06g069330) belongs to the PINOID family and modulates PIN polarity to guide auxin fluxes and direct root growth (Dhonukshe et al., 2015). We further identified 2 LRR containing proteins, including RLK (Solyc07g054120; EXCESS MICROSPOROCYTES1 (EMS1)-like, ITAG4.0), potentially involved in BR signaling. These SNP associations indicate that the activation of signaling cascades in roots is important to evoke waterlogging responses, possibly translating to epinastic bending.

### Hormonal crosstalk during waterlogging-induced epinasty

In the past, it has been shown that waterlogging activates an array of short and long distance hormonal signaling cascades. In our extended set of GWAS targets, we found several genes directly or indirectly involved in hormone biosynthesis and signaling pathways of ethylene, IAA, cytokinin (CK) and abscisic acid (ABA) (Supplemental Table S12). Ethylene, originating from root-borne ACC transport, is the master regulator of the epinastic response during waterlogging (English et al., 1995; Jackson & Campbell, 1976). During the early phase of epinastic bending, our analysis detected a SNP in the ethylene signaling gene Ethylene responsive transcription factor 2A (ERF2A; Solyc07g054220), exceeding the suggestive P-value for leaf 3. Downstream of ethylene, asymmetric growth in the petiole is likely mediated by local auxin redistribution (Lee et al., 2008). The homolog of AUXIN RESPONSE FACTOR 8A (ARF8A; Solyc03g031970), is controlled by the auxin/indole-3-acetic acid gene *SlIAA3/SHY2*, which is known to directly regulate epinastic bending in tomato treated with ethylene (Chaabouni et al., 2009b). While the role of IAA3 in signal polarity seems to be developmentally encoded (Koyama et al., 2010), both AtIAA3/SHY2 and AtARF8 also respond to multiple hormonal cues (Brenner et al., 2005; Chaabouni et al., 2009b). CK for example, directly downregulates *AtARF8* expression in Arabidopsis seedlings (Brenner et al., 2005). In addition, CK and BR signaling, together with changes in *SlIAA3* expression seem to be intertwined in regulating leaf posture in tomato (Li et al., 2016; Xia et al., 2021). While the role of ABA as regulator of stress responses, for example during waterlogging, has been described before (De Ollas et al., 2021), its impact on waterlogging-induced epinasty is unknown. Changes in ABA content have been previously linked with elongation in flooded *R. palustris* (Benschop et al., 2007) and rice (*Oryza sativa*) (Saika et al., 2007).

Besides the well-known classical phytohormones, other signaling compounds with hormone-like properties are gaining more attention. We identified a number of genes in our GWAS analysis which are involved in polyamine (PA) (Amine oxidase family protein Solyc03g031880; Ornithine carbamoyltransferase Solyc12g089210) and melatonin (Aromatic amino acid decarboxylase Solyc07g054280) biosynthesis. Both PA and melatonin have been shown to regulate plant growth and development and abiotic stress responses (reviewed in Chen et al. (2019) and Hardeland (2016)), making them potential mediators of waterlogging responses (Hurng et al., 1994). In tomato, PA metabolism and ethylene biosynthesis are closely intertwined, as both pathways use the same precursor S-adenosyl-L-methionine (Bellés et al., 1992; Takács et al., 2021; Van de Poel et al., 2013).

### Light and hormonal signaling converge during waterlogging

Adequate and fast responses to stress conditions do not always require dedicated signaling pathways to enable survival. For example, plants repurpose the photomorphogenic machinery to activate escape strategies during flooding (van Veen et al., 2013). It is known that AtIAA3 (and AtARF8) combines light and auxin signaling to establish growth (Mao et al., 2020; Xi et al., 2021) in Arabidopsis, while SlIAA3 regulates ethylene-induced leaf bending in tomato (Chaabouni et al., 2009b) and is upregulated during waterlogging (De Ollas et al., 2021). We found SNPs in at least three additional targets, known to operate on the interface between light and hormone signaling (Light-dependent short hypocotyls 1 (LSH1) Solyc02g069510; a GH3 family protein Solyc07g054580, homolog to DWARF IN LIGHT 2 (AtDFL2); Nitric oxide synthase interacting protein, homolog to AtCSU1). In Arabidopsis LSH1 (Zhao et al., 2004) and DFL2 (Takase et al., 2003) integrate red and blue light signals to regulate seedling growth, while CSU1 targets CONSTITUTIVE PHOTOMORPHOGENIC1 (COP1) in dark-grown seedlings (Xu et al., 2014). The possible role of these targets in regulating growth or responses such as epinasty during waterlogging is unknown.

### Petiole polarity as a regulator of asymmetric responses

The involvement of signaling pathways in regulating leaf epinasty eventually require some level of polarity to direct asymmetric growth of the petiole to facilitate downwards bending. At least two target genes (LBD4, Solyc02g069440; Homeobox-leucine zipper protein AtHB14, Solyc03g031760) derived from our GWAS could be related to this process. AtLBD4 is a member of a family of LOB-domain containing proteins, specifically expressed at organ boundaries (Shuai et al., 2002). AtLBD4 itself seems to act at the phloem-procambium interface to regulate vascular development (Smit et al., 2020) and secondary growth in roots downstream of CK signaling (Ye et al., 2021). In *Medicago truncatula*, another LBD transcription factor, ELONGATED PETIOLULE 1 (ELP1), has been shown to define motor organ identity and thus regulate leaf movement (Chen et al., 2012).

Also the formation of the dorsoventral axis during early leaf development is tightly controlled by leaf polarity. The homeobox-leucine zipper protein (Solyc03g031760) identified in our study has not yet been properly annotated nor classified. However, it has resemblance with HOMEODOMAIN GLABROUS 2 (HDG2)-like or MERISTEM LAYER 1 (AtML1) in Arabidopsis, involved in epidermal differentiation. In tomato, it seems to coincide with the LANATA locus, responsible for trichome development (Xie et al., 2022).

## Acknowledgements

We thank the KU Leuven Greenhouse Core Facility for assistance in plant cultivation. This work was funded by the Research Foundation Flanders with a FWO PhD fellowship (11C4319N; 1150822N) to BG and JP respectively, and a FWO research grant (G092419N) to BVdP; by a KU Leuven research grant (C14/18/056) to BVdP and a PhD-back-up grant (DB/17/007/BM) to BG. This work was also established in the framework of the RoxyCost action of the EU (CA18210).

## Author contribution

BG and BVdP designed the experiments, JP and BG phenotyped the tomato accessions, BG analyzed the data, BG performed qPCR analysis and BG and BVdP wrote the manuscript. All authors read and approved the manuscript.

## Supplementary material

Supplemental Figure S1: Growth of 52 tomato accessions after a 3-day waterlogging treatment and subsequent 3-day recovery.

Supplemental Figure S2: Effect of initial plant height on growth of 52 tomato accessions during waterlogging.

Supplemental Figure S3: Natural variation of the effect of waterlogging on 14 different angle descriptors (see Figure 3B) of leaf number 5.

Supplemental Figure S4: Venn diagram showing overlap of annotated genes with (A) suggestive and (B) significant SNPs associated with leaf angle differences between waterlogged and control plants during the early (< 12 h; purple) and late (< 72 h; green) waterlogging phase and during the recovery phase (yellow).

Supplemental Table S1: Phenotypical traits used for the GWAS analyses

Supplemental Table S2: Primers used for RT-qPCR

Supplemental Table S3: GWAS targets related to growth during waterlogging and recovery

Supplemental Table S4: GWAS targets related leaf angle traits and time-series during waterlogging and recovery

Supplemental Table S5: GWAS targets related to early angle responses

Supplemental Table S6: GWAS targets related to early angle responses and recovery

Supplemental Table S7: GWAS targets related to late angle responses

Supplemental Table S8: GWAS targets related to late angle responses and recovery

Supplemental Table S9: GWAS targets related to angle responses during recovery

Supplemental Table S10: GWAS targets related to energy metabolism

Supplemental Table S11: GWAS targets related to DNA, RNA processing

Supplemental Table S12: GWAS targets related to hormone biosynthesis and signaling

Supplemental Table S13: GWAS targets related to light signaling

