## Supplementary Material for "From a different angle: genetic diversity underlies differentiation of waterlogging-induced epinasty in tomato"

### Supplemental material

Supplemental Figure S1: Growth of 52 tomato accessions after a 3-day waterlogging treatment and subsequent 3-day recovery.

Supplemental Figure S2: Effect of initial plant height on growth of 52 tomato accessions during waterlogging (intercept = 0.50442, slope = 0.01975, $R^2_{adj}$ = 0.08, p-value = 0.026).

Supplemental Figure S3: Natural variation of the effect of waterlogging on 14 different angle descriptors (see Figure 3B) of leaf number 5.

Supplemental Figure S4: Venn diagram of annotated genes discovered during different phases of waterlogging-induced epinasty. Diagrams show the overlap of annotated genes with (A) suggestive and (B) significant SNPs associated with leaf angle differences between waterlogged and control plants during the early (< 12 h; purple) and late (< 72 h; green) waterlogging phase and during the recovery phase (yellow).

Supplemental Table S1: Phenotypical traits used for the GWAS analyses

Supplemental Table S2: Primers used for RT-qPCR

Supplemental Table S3: GWAS targets related to growth during waterlogging and recovery

Supplemental Table S4: GWAS targets related leaf angle traits and time-series during waterlogging and recovery

Supplemental Table S5: GWAS targets related to early angle responses

Supplemental Table S6: GWAS targets related to early angle responses and recovery

Supplemental Table S7: GWAS targets related to late angle responses

Supplemental Table S8: GWAS targets related to late angle responses and recovery

Supplemental Table S9: GWAS targets related to angle responses during recovery

Supplemental Table S10: GWAS targets related to energy metabolism

Supplemental Table S11: GWAS targets related to DNA, RNA processing

Supplemental Table S12: GWAS targets related to hormone biosynthesis and signaling

Supplemental Table S13: GWAS targets related to light signaling

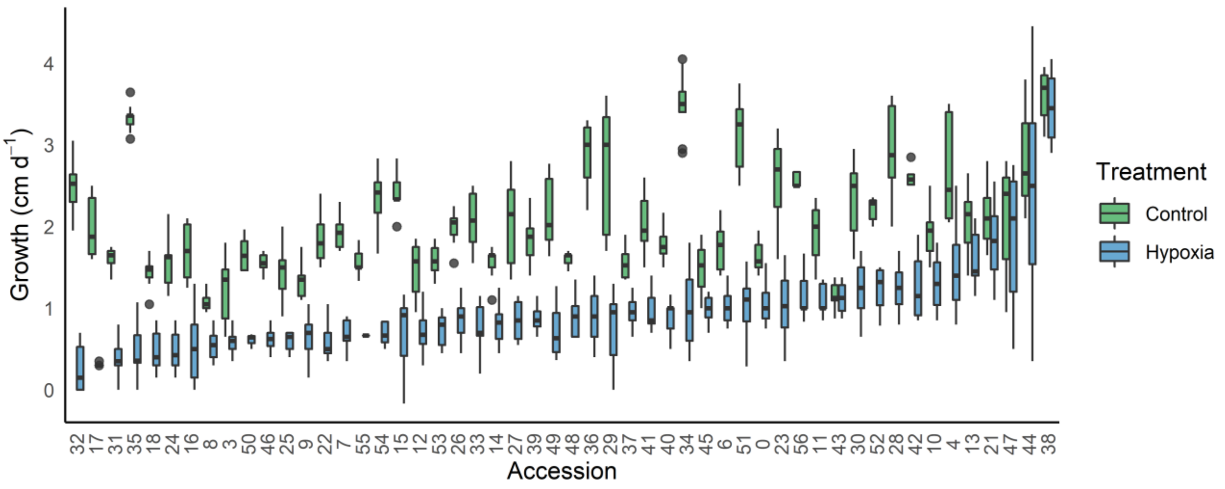


Supplemental Figure S1: Growth of 52 tomato accessions after a 3-day waterlogging treatment and subsequent 3-day recovery.

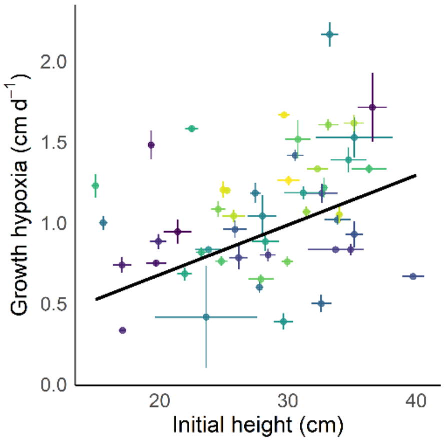


Supplemental Figure S2: Effect of initial plant height on growth of 52 tomato accessions during waterlogging (intercept = 0.50442, slope = 0.01975, $R^2_{adj}$ = 0.08, p-value = 0.026).

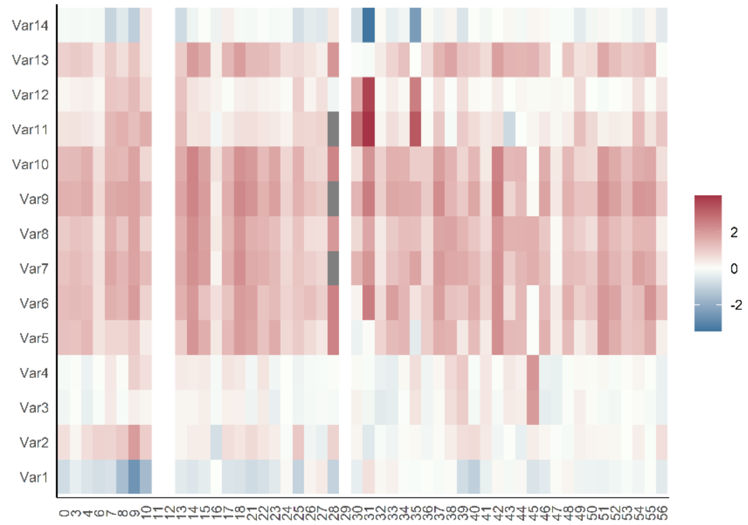


Supplemental Figure S3: Natural variation of the effect of waterlogging on 14 different angle descriptors (see Figure 3B) of leaf number 5. The effect was determined using one-way ANOVA on normalized variable levels.

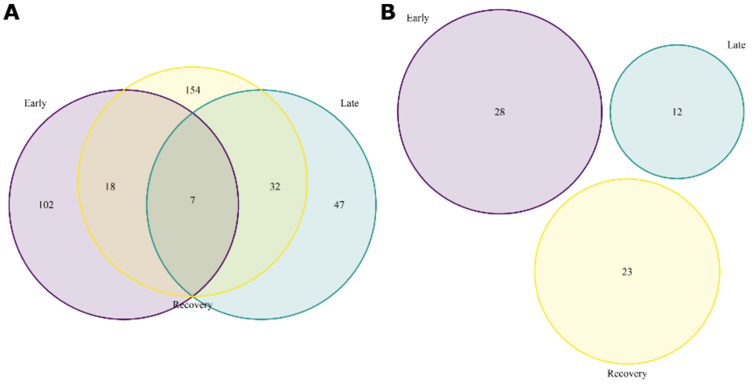


Supplemental Figure S4: Venn diagram of annotated genes discovered during different phases of waterlogging-induced epinasty. Diagrams show the overlap of annotated genes with (A) suggestive and (B) significant SNPs associated with leaf angle differences between waterlogged and control plants during the early (< 12 h; purple) and late (< 72 h; green) waterlogging phase and during the recovery phase (yellow).
